# Performance effects from different shutdown methods of three electrode materials for the power-to-gas application with electromethanogenesis

**DOI:** 10.1101/2024.05.22.595300

**Authors:** Nils Rohbohm, Maren Lang, Johannes Erben, Kurt Gemeinhardt, Nitant Patel, Ivan K. Ilic, Doris Hafenbradl, Jose Rodrigo Quejigo, Largus T. Angenent

## Abstract

Industrial applications of microbial electrochemical systems will require regular maintenance shutdowns, involving inspections and component replacements to extend the lifespan of the system. Here, we examined the impact of such shutdowns on the performance of three electrode materials (*i.e*., platinized titanium, graphite, and nickel) as cathodes in a microbial electrochemical system that would be used for electromethanogenesis in power-to-gas applications. We focused on methane (CH_4_) production from hydrogen (H_2_) and carbon dioxide (CO_2_) using *Methanothermobacter thermautotrophicus*. We showed that the platinized titanium cathode resulted in high volumetric CH_4_ production rates and Coulombic efficiencies. Using a graphite cathode would be more cost-effective than using the platinized titanium cathode in microbial electrochemical systems but showed an inferior performance. The microbial electrochemical system with the nickel cathode showed improvements compared to the graphite cathode. Additionally, this system with a nickel cathode demonstrated the fastest recovery during a shutdown experiment compared to the other two cathodes. Fluctuations in pH and nickel concentrations in the catholyte during power interruptions affected CH_4_ production recovery in the system with the nickel cathode. This research enhances understanding of the integration of biological and electrochemical processes in microbial electrochemical systems, providing insights into electrode selection and operating strategies for effective and sustainable CH_4_ production.

## Introduction

The power-to-gas concept, involving hydrogen (H_2_) or methane (CH_4_), efficiently converts surplus electrical energy into storable gas, facilitating the storage of excess electrical energy in transportable chemical-energy carriers.^[1]^ Electrochaea GmbH has adopted this concept with biomethanation, harnessing excess electrical energy and CO_2_ to generate CH_4_ using *Methanothermobacter thermautotrophicus*, which is explained in detail by Martin and coworkers.^[2]^ While it is possible to achieve high volumetric CH_4_ production rates^[3]^, the process faces challenges, which it has in common with other green methanation processes, due to the high capital costs of conventional water electrolysis. This underscores the pressing need for an alternative method to realize power-to-CH_4_ conversion more effectively.^[4]^ Microbial electrochemistry is an interdisciplinary field that combines principles from biology and electrochemistry. It explores the ability of living microbes to interact with electrodes and utilize electrical energy for various processes. Microbial electrochemical systems have garnered considerable attention due to their potential applications in diverse fields, including energy generation^[5]^, environmental remediation^[6]^, and biotechnology.^[7]^ Microbial electrosynthesis has emerged as a promising approach for sustainable chemical production within microbial electrochemistry. Microbial electrosynthesis involves the utilization of microbes to catalyze chemical reactions by accepting electrons mostly in the form of H_2_ from an cathode.^[8]^ These electrons are then used to drive desired metabolic pathways, including the reduction of CO_2_ to valuable compounds. By harnessing this unique capability, microbial electrosynthesis enables the conversion of CO_2_, which is a greenhouse gas, into valuable chemicals and fuels, providing a potential solution for environmental and economic challenges. Ideally, all the H_2_ produced at the cathode is transformed into CH_4_, generating a fully grid-compatible gas. Furthermore, it could reduce capital expenditures by integrating biology into electrochemical cells. However, microbial electrosynthesis also faces certain limitations and technical obstacles. These challenges include low product yields, inefficient electron transfer, a limited range of products, and the need for a deeper understanding of microbial physiology.^[9]^ Overcoming these hurdles necessitates interdisciplinary collaborations, combining bioelectrochemistry, microbiology, and engineering expertise to develop efficient and economically viable microbial electrosynthesis technologies.

Current density is pivotal, serving as a crucial metric that directly influences the efficiency and effectiveness of microbial electrochemical systems processes. It quantifies the amount of electric current passing through a unit area of the surface of an electrode, thus, dictating reaction rates, product selectivity, and overall performance. Enhancing current densities will result in improved reaction kinetics and yields. Recent studies have explored various approaches to enhance microbial electrosynthesis performance, focusing on either the cathode^[10]^ or the anode^[11]^, as well as optimizing cell configurations.^[12, 13, 14]^ A crucial parameter in establishing a system is ensuring practical energy efficiencies (EEs). Achieving high current densities at low operating cell potentials is essential for practical EEs. Rad et al. demonstrated considerable progress with a hybrid microbial electrosynthesis system, achieving 42% EE during 160 days and a maximum CH_4_ production rate of 669 L(CH_4_) m^-2^ d^-1^ with an average current density of 30 mA cm^-2^ at 2.2 V.^[14]^ However, it was not well shown how much the porous Teflon membrane influenced the performance of the electrochemical cell. In addition, because the microbial community and fermentation broth (referred to as the catholyte) were separated completely from the cathode, it is not clear whether there is a real advantage to integrating electrochemistry and biology into one system.

None of these studies have explored the challenges posed by industrial operating conditions, where operating expenditures (OPEX) and capital expenditures (CAPEX) considerably affect the feasibility of a technology.^[15]^ Embracing industrial conditions is crucial for replicating real-world settings, enabling rigorous testing and validation of processes, products, and technologies. Rovira-Alsina and coworkers used an H-type cell to simulate industrial off-gases with 14% CO_2_ and 12% O_2_ and showed acetate production parity with pure CO_2_ gas.^[16]^ Nonetheless, limitations, such as non-continuous gas feed and open culture conditions, may have impacted microbe survival. In another study, Deutzmann et al. demonstrated an effective microbial electrosynthesis that was powered by intermittent electricity, mirroring a solar energy grid’s power profile.^[17]^ They achieved a >96% Coulombic efficiency (CE) for CH_4_ production using *Methanococcus maripaludis*, showing the compatibility of microbial electrochemical systems with renewable energy fluctuations.

Here, we report on a microbial electrochemical system for simulated maintenance-induced intermittent shutdowns. We tested three types of electrodes, which were platinized titanium, graphite, and nickel, for scale-up processes relevant in terms of availability and cost-effectiveness. Platinized titanium electrodes strike a balance between efficiency and cost. By depositing a thin layer of platinum on titanium, these electrodes retain the catalytic prowess of platinum while mitigating the cost concerns associated with using pure platinum electrodes.^[18]^ This compromise makes platinized titanium electrodes a practical choice and served as the control electrode for our study. Graphite is economical and readily available, making it an attractive option for cost-conscious projects. Similarly, nickel, which is widely recognized for its role in abiotic H_2_ electrosynthesis, presents an additional catalytic effect compared to graphite. We investigated how shutdown methods affect electrode performance. All three electrodes were tested by comparing their performance in producing efficiently CH_4_ from H_2_ and CO_2_ using *M. thermautotrophicus*.

## Materials and methods

### Chemicals, strain, and medium

All chemicals were used as received unless otherwise noted in the text. These chemicals were of technical or analytical grade. The used strain is an adapted laboratory strain of *M. thermautotrophicus* from Martin and coworkers.^[2]^ The medium was a slightly modified version of Martin et al. (2013). The inoculum for the microbial electrochemical system was taken from a CH_4_-producing continuous stirred-tank bioreactor.

### Analytics

Sampling of the microbial electrochemical system occurred twice daily. Evolved gases from the reservoirs were collected in gas bags, and their composition was determined using a gas chromatograph (Agilent micro GC 490, Agilent Technologies, Sanata Clara, California, USA). The gas chromatograph was equipped with two molsieve 5A columns and one pora PLOT U column (Agilent Technologies, USA). A thermal conductivity detector measured N_2_, O_2_, CH_4_, H_2_, and CO_2_. Argon was the carrier gas for one molsieve 5A column, while He was the carrier gas for the other two columns. Trace and major element analysis were conducted using an ion-coupled plasma mass spectrometer (Agilent 7900) with 1% HNO_3_ as the matrix. The samples were centrifuged for 5 min at 25001 relative centrifugal force (Centrifuge 5427 R, Eppendorf; Hamburg, Germany). Every sample was diluted 1:100 in 1% HNO_3_.

All scanning electron microscopy (SEM) images were taken in secondary-electron mode using a Zeiss Crossbeam 550L focused ion beam SEM (Zeiss, Wetzlar, Germany), which was operated with an acceleration voltage of 2 kV. The SEM was equipped with an Oxford Instrument energy dispersive spectrometry detector (Ultim Max, Oxford Instrument, Abingdon, United Kingdom). Elemental maps of iron and carbon were obtained along with corresponding spot analysis. The Nafion 117 samples were coated with a 10-nm deposition of platinum using a BAL-TEC™ SCD 005 sputter coater (Leica Biosystems, Nussloch, Germany). The purpose of the coating was to reduce any charging effects during SEM analysis.

#### Operating conditions

The electrochemical cell that we employed was a two-chamber flow cell from Electrocell (**Figure 1**, Electro MP Cell, Electrocell, Skjern, Denmark). This cell was divided into cathodic and anodic compartments, separated by a proton exchange membrane (**Figure** 1, Nafion 117, FuelCellsEtc, College Station, TX). The cathode (working electrode) materials included platinized titanium (Electrocell, Skjern, Denmark), flat graphite (SGL Carbon, Wiesbaden, Germany), and nickel (Electrocell, Skjern, Denmark), while the anode (counter electrode) material was iridium oxide (16 g m^-2^) on titanium (Electrocell, Skjern, Denmark). The total surface area of the membrane, cathode, and anode was 100 cm^2^. Both compartments were connected to glass reservoirs *via* tubing (Cole-Parmer Instrument Company, Vernon Hill, IL). The cathodic compartment was filled with the medium during experiments, and the anodic compartment contained Milli-Q water. Catholyte and anolyte were recirculated between the glass reservoirs and the electrochemical cell using two peristaltic pumps (Masterflex, Cole-Parmer Instrument Company).

**Figure 1:**
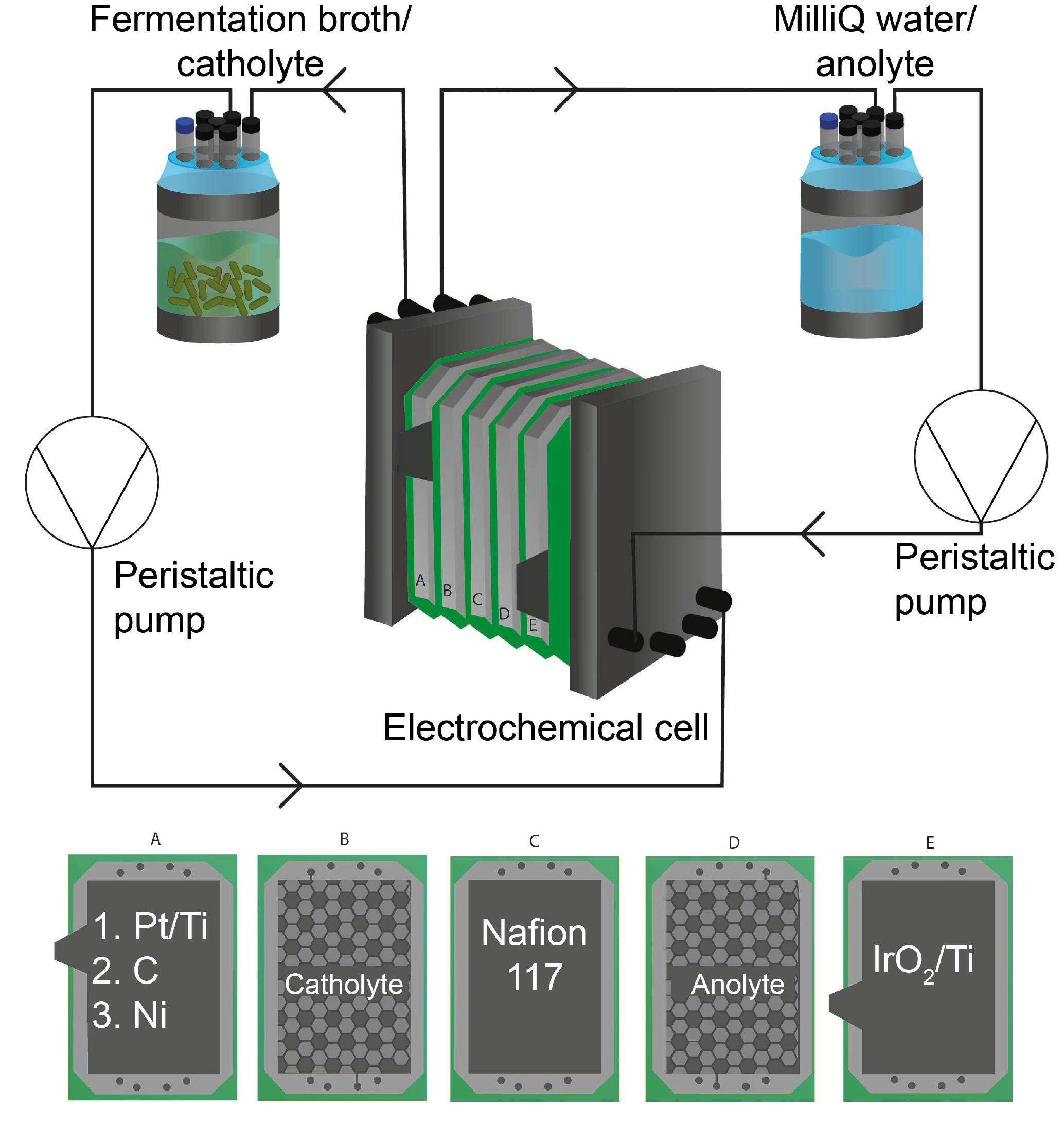
Schematic of the microbial electrochemical system. The catholyte and anolyte recirculation glass vessels are recirculated with peristaltic pumps to the electrochemical cell. The electrochemical cell was separated in five sections with: (A) platinized titanium, graphite, and nickel cathodes; (B) the catholyte flow chamber; (C) the ion-exchange membrane Nafion 117; (D) the anolyte flow chamber; and (E) the iridium oxide on titanium (IrO_2_)anode.

The glass reservoirs featured several openings. One opening served as outgas and was connected to a collection bag (Supel™ Inert foil gas sampling bag, Supelco, Pennsylvania, USA). A tube filled with silica beads preceded the gas bag to dry the off-gas. The pH was measured with a handheld pH meter (Mettler Toledo, Columbus, Ohio). The oxidation-reduction potential (ORP) and conductivity measurements were performed using an ORP probe (Hanna Instruments, Vöhringen, Germany) and a conductivity sensor (Endress + Hauser, Reinach, Switzerland). OD_650_ was measured through a photometer (Lovibond MD 610, Tintometer GmbH, Dortmund, Germany). A water jacket and a recirculating water thermostat (KISS 104A, Huber, Raleigh, NC) were used to heat the water jacket of the reservoirs to 62°C. The temperature distribution was assumed to be uniform due to the high electrolyte recirculation between the glass reservoirs and the electrochemical cell. The total and wet volume of both cathodic and anodic compartments, including glass reservoirs and tubing, was 350 mL for each compartment. The catholyte volume inside the electrochemical cell was 60 mL.

The cathode and anode were connected to a power supply (EA-PS 3040-10 C, EA-Elektroautomatik, Viersen, Germany) with a preset potential of 3V as a two-electrode setup. The current was recorded using a multimeter (Agilent 34972A, Agilent Technologies) with data acquisition through Labview (National Instruments, Austin, Texas). The CO_2_ flow rate was controlled using a mass flow controller (EL-Flow Prestige, Bronkhorst, Kamen, Germany) and adjusted according to the current generated in the electrochemical cell using Labview. A dampening system (R1016823S4000000, Varem, Bovalenta, Italy) was installed downstream of the pump and upstream of the electrochemical cell to reduce pressure fluctuations caused by the peristaltic pump during electrolyte recirculation. For abiotic electrochemical experiments, a similar setup was employed as described above. A reference electrode (3M Ag/AgCl) was introduced into the cathode chamber for the three-electrode setup. The cathode, anode, and reference electrode were connected to a potentiostat (VSP, Bio-logic, Claix, France) that was controlled *via* EC-lab software (Bio-logic).

### Startup and shutdown procedure

When starting the electrochemical cell, the power source was set to 3 V, the water thermostat was adjusted to achieve a temperature of 62°C within the glass reservoirs, and the recirculation pump was set to 120 ml min^-1^. OD_650_, conductivity, temperature, and pH of the catholyte and anolyte were analyzed daily by sampling 10 ml. Initially, the cell was inoculated with a starting OD_650_ of 1.

Each experiment was run for multiple weeks, one week comprising 5 days of operation followed by 2 days of shutdown. Catholyte and anolyte levels were adjusted on the second and the fourth day during the platinized titanium experiment and on the third day of every week during the graphite and nickel experiments. The anolyte was refilled with distilled water, and the catholyte was removed to reach the original liquid volumes. A concentrated medium was added to compensate for the loss of nutrients due to catholyte removal and sampling. The setup was started at the beginning of each week by starting the heating and the peristaltic pump. Once the operating temperature was reached, the power source was set to 3V, and the CO_2_ supply started. During the nickel experiment, liquid samples for trace element analysis were taken right before switching on the power source, 2h after switching on the power source, and on Day 3 and 5 of each week. On Day 5, the system was shut down with the following procedure: the heating and power source were switched off, the electrical circuit between the power source and the cell was disconnected (open-circuit mode), and the CO_2_ supply was stopped. Once the temperature in the reservoirs decreased to < 40°C, the pump was shut off. During the experiments with Nickel cathode, the shutdown procedure was altered in some weeks where the cell was polarized at 0.9V or short-circuited.

### Equations

The Coulombic efficiency (CE) was calculated according to Kracke and colleagues ^[10]^:

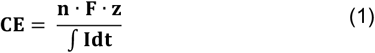

Where n is the molar quantity of produced CH_4_ and H_2_, F the Faraday constant, z the number of electrons to form CH_4_ or H_2_, and I the current of the microbial electrochemical system. The EE was calculated as follows ^[19]^:

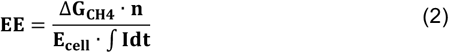

Where ΔG_CH4_ is the Gibbs free energy of CH_4_ (890.4 kJ mol^-1^) and E_cell_ the applied cell voltage (3 V).

## Results and Discussion

### The microbial electrochemical system using a platinized titanium cathode exhibited higher production rates compared to those with graphite and nickel cathodes during experiments with repeated interrupted power supply

Three-electrode types were run in a microbial electrochemical system with defined electrolyte flow rates (120 mL min^-1^). Given the importance of pH control as a cost determinant, pH monitoring was undertaken without concurrent pH regulation. The overarching concept was to establish an equilibrium state between catholyte and anolyte pH, obviating the need for bidirectional pH control measures even if a pH gradient would be created. At first, the experiment ran for three weeks for each of the three electrode materials with consecutive 5 days of runtime and 2 days without current. During the two power supply interruptions, the microbial electrochemical cell was disconnected from the power source (open-circuit).

During the comparative analysis of cathode materials, platinized titanium, serving as a benchmark for this study, demonstrated a higher volumetric CH_4_ production rate and a higher CE to CH_4_. The microbial electrochemical system with this cathode achieved a peak volumetric CH_4_ production rate of 16.9 L L^−1^ d^−1^ (volume of CH_4_ per microbial electrochemical system cell volume per day) and a CH_4_ Coulombic efficiency (CE-CH_4_) of 92.6% (**Figure 2A**). This relatively high volumetric production rate can be attributed to its excellent catalytic properties of platinum, which increased the overall current compared to other materials (**Figure 2**).^[18]^ During the first week, the production rate steadily increased with the increasing current during the starting period, reaching a maximum of 16.8 L L^−1^ d^−1^ at 90.2% CH_4_-CE (**Figure 2A**). Following the first shutdown, the production rate progressively climbed, peaking at 16.9 L L^−1^ d^−1^ by the end of the second week. The microbes responded better to the interruptions after the second shutdown, achieving a volumetric CH_4_ production rate of 16.7 L L^−1^ d^−1^ within two days (**Figure 2A**).

**Figure 2:**
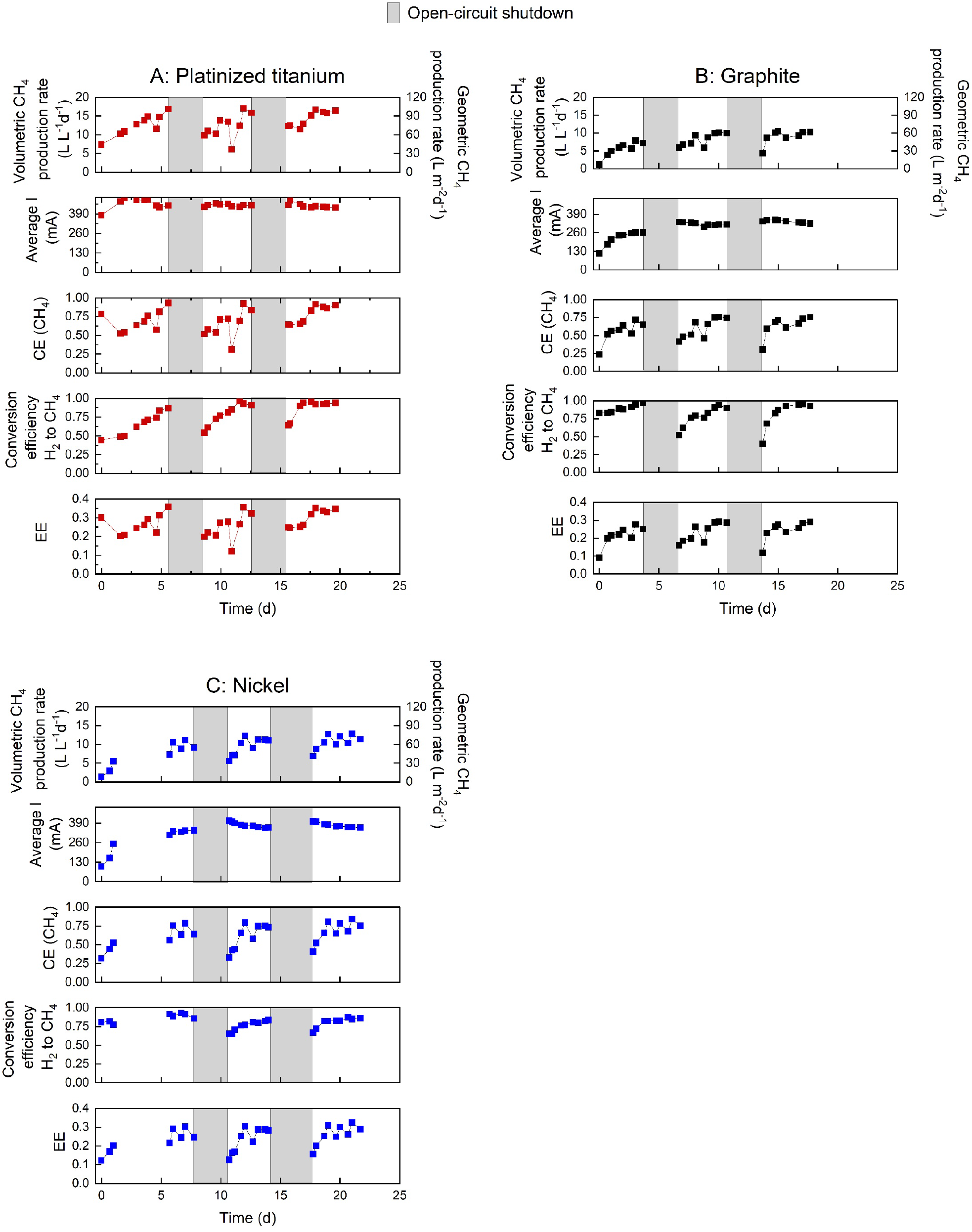
Comparison of different electrode materials during a 5-day operation periods followed by 2-day shutdown periods with: (A) platinized titanium; (B) graphite; and (C) nickel cathodes. Shown are for each cathode: the volumetric CH_4_ production rate (Y1 Axis top); the geometric production rate (Y2-axis top graph); average current (I) of a sample period; the Coulombic efficiency (CE) of CH_4_, the conversion efficiency of H_2_ to CH_4_; and the energy efficiency (EE).

The microbial electrochemical system using graphite as cathode, while cost-effective and accessible, underperformed in comparison to the platinized titanium cathode. The microbial electrochemical system with the graphite cathode achieved a lower maximum volumetric CH_4_ production rate of 10.4 L L^−1^ d^−1^ and CH_4_-CE of 71.8% (**Figure 2B**). The lower production rate can be attributed to a lower current because of the absence of a catalyst.

The graphite cathode exhibited its highest average current after the second shutdown with 350 mA, while the platinized titanium cathode generated an average current of 498 mA, which is 1.4 fold higher (**Figure 2A and B**). Initially, the average current for the graphite cathode was 264 mA at the end of the first week (**Figure 2A**). After the first shutdown, the average current rose from 264 mA to 336 mA, reaching a maximum volumetric CH_4_ production rate of 10.1 L L^−1^ d^−1^ with a CH_4_-CE of 75.8%. After the second shutdown, the volumetric CH_4_ production rate stabilized at a maximum of 10.5 L L^−1^ d^−1^, but the CH_4_-CE never surpassed 75.5% (**Figure 2B**), which is inferior when compared to the >90% CH_4_-CE of the platinized titanium cathode (**Figure 2A**).

The microbial electrochemical system with the nickel cathode showed improvement compared to the graphite cathode in terms of average current, which is attributed to a higher conductivity and catalytic activity (**Figure 2C**). The average current reached a maximum of 405 mA, which is 15.5% higher than the graphite cathode and 18.5% lower than the platinized titanium cathode (**Figure 2**). The microbial electrochemical system with the nickel cathode had a maximum volumetric CH_4_ production rate and CH_4_-CE of 12.8 L L^−1^ d^−1^ and 84.4%, respectively (**Figure 2C**). However, it fell short of matching the performance of the microbial electrochemical system with a platinized titanium cathode. After an initial 4-day current ramp-up, the microbial electrochemical system with a nickel cathode reached stable volumetric CH_4_ production rates at 11.2 L L^−1^ d^−1^ and 78.8% CH_4_-CE (**Figure 2C**). After the first shutdown, it took only a day to return to pre-shutdown production levels, achieving a maximum of 12.3 L L^−1^ d^−1^ with a CH_4_-CE of 79.4%. A similar pattern was observed in the third week, peaking at 12.8 L L^−1^ d^−1^ and 84.4% CH_4_-CE (**Figure 2C**).

Another critical factor for industrial readiness is energy efficiency (EE). In this study, the average EEs for microbial electrochemical systems with platinized titanium, graphite, and nickel cathodes were 23.7%, 22.2%, and 21.2%, respectively, with their maximum efficiencies reaching 35.8%, 29.2%, and 32.4%, respectively (**Figure 2**). These values align closely with existing literature, showing similar current densities and cell voltages.^[12, 20, 21]^ When comparing platinized titanium, graphite, and nickel cathodes (**Figure 2**), it is apparent that each material has its unique strengths and weaknesses. However, the platinized titanium cathode consistently surpassed the performance of the other cathodes in this study. The reason for the overall lower CH_4_-CE of the graphite and nickel cathodes during the microbial electrochemical system experiments was the lower H_2_ production or the H_2_ loss through a suboptimal system. The average CEs of H_2_ were 80.4% and 69.8% for the graphite and the nickel cathodes in the microbial electrochemical systems, respectively (**Figure S1**). The microbial electrochemical system with a platinized titanium cathode achieved an H_2_-CE of 91.3%, showing a higher availability of H_2_ in the system and a higher efficiency in producing H_2_ (**Figure S1**). A better comparison would be the conversion efficiency of H_2_ to CH_4_. Because H_2_-CE never consistently reached 100%, the CH_4_-CE is constrained due to the calculation with the total number of electrons transferred in the electrochemical reaction. Conversely, the efficiency of converting H_2_ to CH_4_ is calculated based on the measured H_2_, thus, circumventing the limitations posed by H_2_ production inefficiencies. Here, the microbial electrochemical system with a platinized titanium cathode showed the lowest conversion efficiency at 77.2% followed by the nickel cathode at 80.6% and the graphite cathode at 83.1% (**Figure 2**). The reduced conversion efficiency observed in platinized titanium is attributed to the lower CH_4_ production rate, despite the high H_2_ production (**Figure S1**). Regardless, despite interruptions in the power supply, the microbial electrochemical system with a platinized titanium cathode consistently showed superior performances, which we attribute to its superior catalytic properties.

### Low pH lead to nickel dissolution and influenced the recovery of CH_4_ production

Deutzmann and colleagues demonstrated that *M. maripaludis* exhibited robust CH_4_ recovery despite varying shutdown times.^[17]^ In scenarios with intermittent power supply for electrochemical CH_4_ production, the choice of electrode material considerably impacts the recovery process. We compared the three cathode materials and assessed which electrode material was able to recover CH_4_ production with an H_2_ to CH_4_ conversion efficiency goal of >80%. The platinized titanium cathode recovered in 1.9 days, the graphite cathode slightly lagged, taking 2.4 days and the nickel cathode showed a rapid recovery response of 1.9 days to reach 80% H_2_ to CH_4_ conversion efficiency. Notably, after the second power shutdown, all three cathodes reached >80% conversion efficiency within 1.0 day, with platinized titanium achieving the highest H_2_ to CH_4_ conversion efficiency (89.9%, **Figure 2A**). Platinized titanium and graphite had similar average H_2_ to CH_4_ conversion efficiencies (93.1% and 90.3%, respectively, **Figure 2A** and **B**), while the nickel cathode averaged 84% (**Figure 2C**).

While operating conditions for all three cathodes remained consistent, a considerable pH difference emerged, especially for the nickel cathode. Initially, pH values for all microbial electrochemical systems ranged from 6.7 to 7.5 in the catholyte (**Figure S2**). However, a shift towards higher pH values occurred for the microbial electrochemical system using platinized titanium and graphite cathodes. In contrast, the microbial electrochemical system with the nickel cathode experienced a pH decrease to 5.9 and 6.3 during the first and second power supply shutdowns. Unfortunately, the specific cause for the pH drop is unknown to us. This pH shift, lowering the pH from the optimal growth range, led to an elevated nickel concentration in the medium, potentially affecting CH_4_ production recovery. The initial nickel content (**Figure S3A**) was 0.29 ppm, and increased to 3.11 ppm before the shutdown, further elevated to 5.15 ppm after the shutdown, and decreased to 2.38 ppm before the second shutdown. After the microbial electrochemical system was reactivated after the second shutdown, the nickel concentration had a modest increase to 3.46 ppm, potentially influencing the observed accelerated recovery in CH_4_ production. In summary, the CH_4_ recovery for all three cathode materials was less than a day after an adaptation to the power shutdowns. The lower conversion efficiency for the microbial electrochemical system with the nickel cathode could be explained by the lower pH rather than higher nickel concentrations in the fermentation broth as we show below.

### Short-circuit shutdowns had a higher impact on CH_4_ production and recovery times compared to polarized and open-circuit shutdowns

Nickel cathodes have the benefit of being durable and stable while having good catalytic properties to produce H_2._ It is, therefore, an ideal candidate for large-scale applications. Additionally, nickel supports the growth of *M. thermautotrophicus* in electromethanogenesis *via* nickel enzymes.^[22, 23]^ Therefore, using nickel cathodes in electromethanogenesis offers a favorable environment for their growth, ensuring that nickel availability does not become a limiting factor. However, similar to many other compounds, an upper concentration limit exists beyond which it becomes toxic. In the preceding section, we demonstrated that a lower pH value in combination with elevated nickel concentrations slowed CH_4_ production recovery, although the differences observed after the second interruption period were relatively minor. To comprehensively assess the impact of nickel concentrations on electromethanogenesis, we extended the experiments for the microbial electrochemical system with the nickel cathode involving repeated power supply interruptions. The experiment was continued following the same experimental procedure. This involved a 5-day run time followed by a 2-day power supply interruption. We explored the potential occurrence of side reactions depending on the electrical connectivity of the microbial electrochemical system. Three interruption modes were employed: **1)** open-circuit mode in which the electrical circuit was interrupted by disconnecting a wire, and which was a continuation of the previous operating conditions; **2)** polarized mode, involving the application of an external potential of 0.9 V; and **3)** short-circuit mode for which the anode and cathode were connected to the power source by wires, but no current was set, allowing electrons to flow freely between cathode and anode (**Figure 3**).

**Figure 3:**
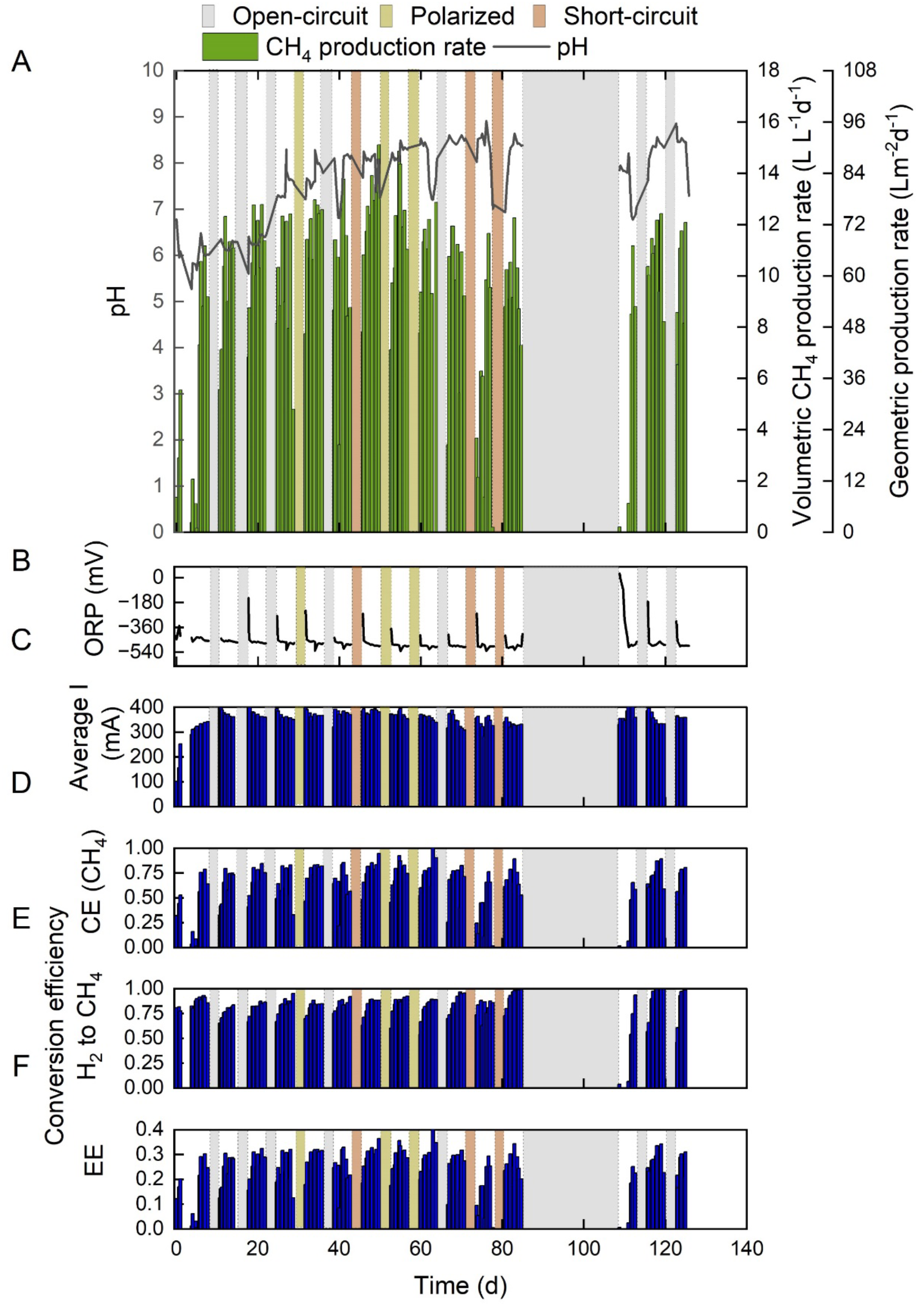
Period of the nickel cathode experiment with different shutdown methods coded in grey open-circuit shutdown, light yellow polarized shutdown, and light red short-circuit shutdown. The different graphs show: (A) the microbial electrochemical system cell volumetric (Y2-axis) and geometric (Y3-axis) CH_4_ production rate as well as the pH; (B) the oxidation-reduction potential (ORP); (C) the average current of a sample period; (D) the Coulombic efficiency (CE) of CH_4;_ (E) the conversion efficiency of H_2_ to CH_4_; and (F) the energy efficiency (EE) of the microbial electrochemical system

Accordingly, we continued the experiment with the nickel cathode after the two open-circuit shutdown periods (**Figure 2C** and **3A**). Following the third open-circuit shutdown, the nickel concentration was 2.13 ppm with a CH_4_ production recovery period of one day and a maximum volumetric CH_4_ production rate at 12.4 L L^-1^ d^-1^ (**Figure S3** and **3A**), which we anticipated based on the second open-circuit shutdown. While polarized and open-circuit shutdowns during weeks 4 and 5 showed no considerable differences in nickel concentrations (3.02 and 2.56 ppm, respectively), a notable increase in the nickel concentration occurred after a short-circuit shutdown during week 6 (7.95 ppm). A higher nickel concentration was reached despite a relatively high pH of 7.86 (**Figure 3A**), which was unanticipated by us. The short-circuit event enhanced nickel dissolution. However, the higher nickel concentration at a relatively high pH achieved the CH_4_ production recovery time to achieve at least 80% CH_4_ conversion efficiency in only half a day (**Figure 3E**). Despite the higher initial nickel concentrations, the microbial electrochemical system reached its highest volumetric CH_4_ production rate of 15.1 L L^-1^ d^-1^ (**Figure 3A**). Therefore, higher nickel concentrations had no negative impact on CH_4_ production because a nickel concentration at which it becomes toxic was not reached. On the contrary, an improved CH_4_ production can be explained by the stimulating effect of Nickel for certain Nickel-enzyme activities. This result shows that the earlier shift to a longer CH_4_ production recovery period was due to the lower pH values because, for the short-circuit mode, the pH had stayed slightly alkaline.

Two polarized and one open-circuit shutdowns followed the short-circuit shutdown through weeks 7 to 9, decreasing nickel concentrations (**Figure S3A**) and restoring nickel concentrations to values before the week 6 short-circuit shutdown. The idea was to further investigate the impact of short-circuit shutdowns on nickel dissolution. Interestingly, H_2_ to CH_4_ conversion efficiencies decreased to levels that we had observed before the first short-circuit shutdown. During week 10, another short-circuit shutdown resulted in a higher nickel concentration (7.36 ppm), but it did not affect the CH_4_ conversion efficiency, while still recovering within half a day. In contrast to the first short-circuit mode shutdown, the volumetric CH_4_ production rates did not exceed other previous shutdown methods (**Figure 3** and **S3A**). While the open-circuit shutdown mode was optimal to avoid excessive nickel concentration increases due to nickel dissolution, all shutdown methods were viable because increased nickel concentrations did not impair CH_4_ conversion efficiency. It is crucial to note that a higher frequency of short-circuit shutdowns could potentially harm the nickel cathode due to excessive nickel dissolution, leading to degradation and a loss of catalytic efficiency in H_2_ production. Overall, utilizing the open-circuit or polarized modes instead of the short-circuit mode is beneficial for the microbial electrochemical system.

### Gradual degradation of the nickel cathode in a microbial electrochemical system throughout a ten-weeks operating period was less impacted by the different shutdown modes than by other factors

We anticipated that the performance of the nickel cathode would gradually decline throughout the operating period due to: **1)** morphology changes of the catalysts; **2)** electrolyte impurities; and **3)** corrosion. During the operating period of 10 weeks for the microbial electrochemical system with a nickel cathode, the average current decreased from 378 mA for week 1 to 334 mA for week 10, corresponding to 11.6% degradation (**Figure 3C**). To further measure the deterioration of the nickel cathode, we assessed the reduction in current throughout the operating period. Therefore, a linear regression was used for each of the restarts of the microbial electrochemical system until the next shutdown occurred. The regression analysis revealed no trend indicating that not a particular shutdown mode degraded the nickel cathode more. The observed current degradation spanned a range from -14.1 mA d^-1^ to -1.1 mA d^-1^ and occurred without any shutdown method impacting the decrease in current throughout time (**Figure S4**). The nickel cathode showed signs of degradation throughout the operating period, but the shutdown modes did not seem to be responsible.

One of the degradation issues that can arise is electroplating at the cathode with metals from the catholyte. This would be seen by the decreased electrolyte concentration of the electroplated compound. To address this concern, we monitored the electrolyte/medium throughout the experiment and assessed the concentration of metal compounds (Mg, Mn, Fe, Co, Zn, Se, and Mo), excluding nickel (refer to **Figure S3**). Among these elements, selenium (Se) displayed no considerable change in concentrations (**Figure S3F**). The detected calcium was not added to the medium; we assume contamination from an unknown source. Similarly, zinc (Zn) was detected, potentially indicating contamination from a reactor component containing zinc (**Figure S3B**).^[24]^ Nevertheless, this contamination did not appear to have a discernible impact on the operation of the microbial electrochemical system.

The most notable effects were observed with magnesium (Mg) and iron (Fe), which showed a near depletion of these metals in the electrolyte (**Figure S3G** and **H**). The decreased Fe concentration could indicate electroplating at the cathode. Intriguingly, despite the importance of iron in iron-sulfur clusters and magnesium in ATP and ADP-dependent reactions^[25]^, this depletion did not appear to impede CH_4_ production. A study reported decreased metal deposition at the cathode when a concentrated trace element solution was used with a chelating agent.^[26]^ In our study, we used nitrilotriacetic acid as the chelating agent, which may have similarly reduced electroplating (**Figure S3**). However, the fermentation broth was saturated with carbonate, potentially leading to the precipitation and subsequent filtration of Mg and Fe during the preparation for ICP-MS measurement. Finally, manganese (Mn), cobalt (Co), and molybdenum (Mo) (**Figure S3C, D**, and **E**) exhibited a sharp initial decrease in their concentration, which later stabilized. The concentration of the elements was measured using cell-free effluent, and the decrease in concentration could also have resulted from *M. thermautotrophicus* taking the elements for nutrition instead of electroplating. Furthermore, once per week, the electrolyte level was adjusted with our medium, which resulted in an immediate increase in the concentration and a slower decrease throughout the operating period of the microbial electrochemical system.

If electroplating was not the clear cause of the decrease in current density, it suggests that other potentially harmful reactions may have occurred. These potential reactions may have occurred at the cathode surface or during the shutdown period, corroding the nickel cathode and decreasing its performance. A degradation possibility occurs during self-discharge and open-circuit voltage, where the nickel cathode may have corroded, leaching either Ni^2+^ or Ni(OH)_2_ into the solution, which we observed after each shutdown method (**Figure S3**). Furthermore, further oxidation to β-Ni(OH)_2_ or NiO could have occurred due to the elevated O_2_ content during the initial phase (**Figures. S5** and **S6**). An abiotic study conducted by Kim and colleagues explored the recyclability of nickel cathodes through successive shutdown and startup cycles of the electrochemical cell, and an increased degradation of the nickel cathode was observed.^[27]^ In our study, the corroded Ni^2+^ could be electroplated at the bare nickel cathode, reducing the reaction sites for the H_2_ evolution reaction. However, substantial proof for these side reactions could not be found. Overall, the microbial electrochemical system should not be interrupted frequently in open-circuit to avoid corrosion. In case of an interruption and leaching of nickel into the electrolyte, any O_2_ contamination should be prevented to avoid further corrosion reactions.

### More prolonged power interruption influences the recovery of CH_4_ production

In a circumstance where a more extended interruption of the system occurs due to a breakdown of components, a speedy recovery of the system is the desired outcome. We showed that after an initial adaption of three consecutive shutdowns, *M. thermautotrophicus* could quickly recover to >80% H_2_ to CH_4_ conversion efficiencies within a day. We then examined if our system could quickly recover its CH_4_ production after a 24-day shutdown. We initiated an open-circuit shutdown mode for this relatively long period (**Figure 3**), and we did not use additional prevention methods to ensure the survivability of the microbe. After restarting the microbial electrochemical system, the H_2_ level showed near 100% H_2_-CE (**Figure S7**); consequently, no CH_4_ was produced. *M. thermautotrophicus* has rapid growth, with a doubling time of 3 h. However, a CH_4_ conversion efficiency of <10% suggests that a recovery or improvement in CH_4_ output is unlikely without intervention or optimization of the process.

A long-term shutdown can increase the likelihood of O_2_ contamination. Even if the microbe could survive long exposures to O_2 [28]_, in our study, the extended shutdown and the increased ORP value are two variables associated to the decrease in the performance. The ORP of the microbial electrochemical system was 32.2 mV at the restart of the microbial electrochemical system at which point no methanogenesis could occur^[29]^ (**Figure 3B**), which indicated O_2_ exposure. The ORP reached -304.2 mV after only 1 day, which is beneficial for methanogenesis. However, the ORP further decreased to -504 mV without *M. thermautotrophicus* being able to produce CH_4_. The microbial electrochemical system was then reinoculated with fresh inoculum from a continuous stirred-tank bioreactor, which reestablished CH_4_ production quickly. The conversion efficiency of H_2_ to CH_4_ of the microbial electrochemical system reached 75% after 1 day and 94% after 1.7 days (**Figure 3E**). The subsequent open-circuit shutdowns did not negatively impact the recovery of CH_4_ (**Figure 3**). For future applications, the glass recirculating vessel should be designed and built to prevent O_2_ leakage, which increased the ORP during our long-term shutdown. The above-mentioned O_2_ problem was due to the relatively small experimental setup and would be less of a problem at industrial scales with stainless steel materials.

### Short-term O_2_ exposure did not harm the CH_4_ recovery time of the microbes

To facilitate the growth of methanogens, the system must maintain anoxic conditions, primarily due to the absence of mechanisms for O_2_ detoxification. Fortunately, *M. thermautotrophicus* exhibits a higher O_2_ exposure tolerance than other methanogens.^[28]^ However, given the setup (customized drying tube for the off-gas), there was O_2_ contamination, which could have impeded the methanation process. During our experiments, we monitored O_2_ concentrations, yet we did not observe a substantial reduction in the methanation process, even with extended exposure to O_2_ (**Figures S5** and **S6**). Notably, high O_2_ concentrations were detected at the beginning and after restart of the experiment with all three cathodes (**Figure S5**). While it took several days to achieve high H_2_ conversion into CH_4_, the O_2_ levels remained consistently low and sometimes exhibited spikes, which, interestingly, did not seem to affect the production rates (**Figures 3** and **S6**). Because the drying tube was placed in the off-gas, O_2_ contamination mainly occurred during the shutdown periods. In the case of the prolonged nickel experiment, high percentages of O_2_ were observed as well, with negligible impact on the conversion efficiency (**Figure 3E**). Because the temperature was switched off during the shutdown periods, the microbes would have been dormant, which might have extended their survival.

However, it is worth noting that in some instances, O_2_ levels spiked (**Figure S6**), adversely affecting the overall production rate and reducing both the EE and CE of H_2_ and CH_4_ (**Figures 3F** and **S7**). When these O_2_ spikes occurred, the O_2_ to N_2_ ratio corresponded to air, suggesting a possible system leak, likely stemming from the customized drying tube (for further explanation, see material and methods), which we connected before the collection bag. Nevertheless, *M. thermautotrophicus* demonstrated its capacity to sustain the electromethanation process even during such leakages. In summary, despite the potential challenges posed by O_2_ leakage and the associated variability in production efficiency, the resilience of *M. thermautotrophicus* and the ability of the system to maintain anoxic conditions underscore the robustness of the electromethanation process, even in the face of unexpected environmental or system fluctuations. However, we also showed that the 24-day O_2_ leakage and exposure should be prevented.

### The operating conditions of the microbial electrochemical cell promote water and gas crossover through the ion-exchange membrane

In general, one of the performance losses in electrochemical cells arises from the ion exchange membrane^[30]^, which poses a challenge that must be addressed for successful implementation in an industrial setting. When electrodes are polarized, three modes of mass transport become relevant: convection, migration, and diffusion. While diffusion through the ion exchange membrane is restricted, mass transport *via* migration and convection can occur when an electric field is applied, and a pressure difference exists between the two chambers. Notably, during the microbial electrochemical system with the nickel cathode experiment, the catholyte volume increased in one week by 53.9 mL on average, while the anolyte volume decreased by 100.7 mL. The reduction in anolyte volume can be attributed to the fast migration of produced H^+^-ions (Grotthuss’ mechanism), leading to a decrease in the overall anolyte volume. These H^+^-ions can cross the cation-exchange membrane and become neutralized in the catholyte, resulting in the dilution of the catholyte. Osmotic pressure is presumably the primary factor in water crossover, where the higher salinity in the cathode chamber compared to the anode chamber drives water to flow from the anode to the cathode. Another plausible explanation is that water transport is driven by hydraulic pressure due to a slight overpressure on the anolyte side in the microbial electrochemical cell. Despite our efforts to minimize the hydraulic overpressure by maintaining identical liquid flow rates in both chambers and using a dampening system, it cannot be ruled out that hydraulic overpressure might have occurred.

The occurrence of water crossover is unsurprising. However, it was observed that CO_2_ also crossed over from the catholyte to the anolyte during operation (**Figure S8**). For an anion-exchange membrane, which we did not use here, CO_2_ crossover occurs due to the formation of carbonate species, which can migrate under the influence of the applied electric field. The presence of CO_2_ in the anode chamber is counterintuitive, because the Nafion 117 membrane, a proton-exchange membrane, should ideally prevent anion crossover. We suspect three potential explanations for this phenomenon: **i)** the precipitation of carbonate species on the membrane; **ii)** hydrophobic property loss of the membrane; **iii)** swelling of the membrane from increased temperature exposure, and **iv)** CO_2_ permeability from the Nafion membrane. Because CO_2_ crossover was evident from the beginning, it is unlikely that the Nafion 117 membrane was obstructed. However, we observed undefined precipitates on the Nafion membrane after the experiment, which we analyzed with SEM and EDX. Suspected carbonate precipitates from continuous CO_2_ feeding could clog the membrane, while the acidic anolyte (monitored, not continuously tracked, pH of 1.6 ± 0.2) may counteract this by dissolving the precipitates, enabling CO_2_ crossover. Simultaneously, the temperature-sensitive swelling of Nafion 117 might further influence carbonate behavior, contributing to both precipitation and transmembrane crossing.^[31, 32]^ Finally, Nafion 117 naturally permeates CO_2_ and the permeability of CO_2_ is increased when the Nafion membrane is hydrated.^[33]^

The SEM analysis of the Nafion 117 membrane, post-experimentation, revealed the formation of precipitates during the microbial electrochemical system operations (**Figure S9**). Compared to an unused Nafion 117 membrane (**Figures S9A** and **B**), the post-experiment Nafion 117, which was taken after the experiment with the microbial electrochemical system with the platinized titanium cathode, exhibited a uniformly distributed layer of precipitate (**Figures S9C** and **D**). Upon closer inspection at higher magnifications, the precipitate appeared amorphous (**Figures S9E** and **F**). EDX spectroscopy analysis identified peaks corresponding to sulfur, fluorine, carbon, phosphorus, and iron within this amorphous precipitate (**Figure S10**). The presence of sulfur and fluorine aligns with the composition of Nafion, while phosphorus and iron are likely constituents of the precipitate. Although carbonate species were not directly detectable, their involvement in the precipitation, driven by substantial CO_2_ crossover, should not be disregarded. Carbonate presence was measured by acidifying a 1 cm^2^ Nafion 117 segment placed in a sealed serum vial, with GC measurements indicating a CO_2_ concentration of 1.9 ± 0.6 mmol cm^-2^, affirming carbonate accumulation on the membrane. Calcinating the Nafion 117 membrane and subsequent ICP-MS analysis of the residue confirmed the detection of all elements present in the medium (**Figure S11**). Furthermore, the abundance of elements that form carbonates (Na, Mg, K, Fe) is higher compared to the elements that do not form carbonates. Despite attempts to mitigate mass transport effects through a dampening system, water and CO_2_ crossover phenomena underscore the challenges in achieving optimal operating efficiency. Future research should address these membrane-related challenges to enhance the industrial applicability of microbial electrochemical technologies.

### The flow rate influences the current generation and has a higher effect on the microbial electrochemical system performance

Resistance considerably influences microbial electrochemical system performance, with solution, charge transfer, and diffusion resistance being the primary factors.^[34, 35]^ These resistances, resulting from ohmic limitations, kinetic constraints, and mass transport phenomena, respectively, can be modulated by adjusting the flow rate of the electrolyte. However, finding the optimal flow rate is crucial, as it is pivotal in removing gas bubbles and maintaining active reaction sites.

To determine the optimal flow rate for our setup, we conducted experiments using graphite cathodes under three different recirculation flow rates: 100 mL min^-1^, 200 mL min^-1^, and 300 mL min^-1^. *Prior* to the experiments, we preconditioned the abiotic electrochemical cell for 20 h at 3 V to ensure uniform conditions across tests. We maintained the anolyte and catholyte pH during the flow rate experiments at 2 and 7, respectively. We employed cyclic voltammetry, sweeping potentials from -0.7 to -1.15 V vs. Ag/AgCl (3M KCl), at a scan rate of 25 mV s^-1^ to assess the impact of flow rates on system performance. The results indicated that the highest current (-250 mA in **Figure 4**) was achieved at the lowest flow rate (100 mL min^-1^), while the lowest current, -156 mA, was observed at the highest flow rate (300 mL min^-1^) at -1.15 V. Interestingly, the intermediate flow rate (200 mL min^-1^) presented a moderate reduction in current density, indicating a potential optimal middle ground. The relationship between flow rate and system efficiency is not linear, as excessively high flow rates can adversely affect efficiency despite reducing resistive losses from bubble formation. Our empirical findings hint at an optimal flow rate below the 4 L min^-1^ maximum recommended by the manufacturer (Electrocell, Denmark), underscoring the importance of tailored operating adjustments based on specific system performance. This suggests that an optimal current generation in our microbial electrochemical system is achieved with a flow rate that is neither too low nor too high, ideally around 100 to 200 mL min^-1^. This observation suggests that previous biotic experiments have been conducted under optimal flow conditions.

**Figure 4:**
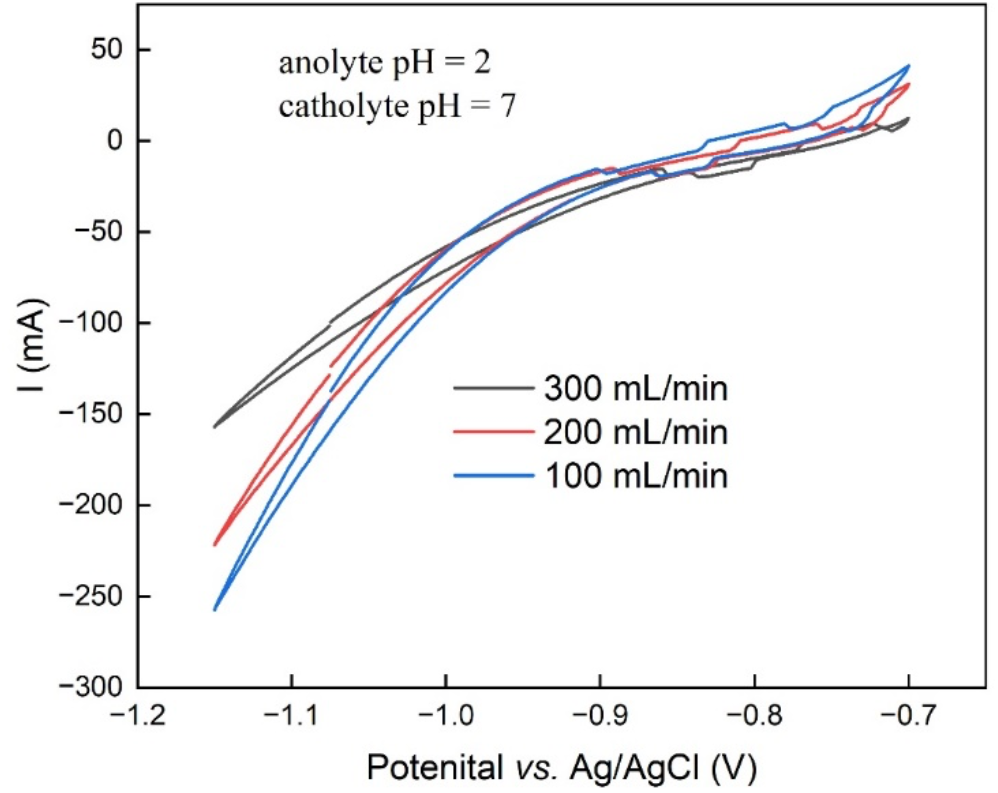
Cyclic voltammograms on the microbial electrochemical system performance influence of flow rate at constant pH. The microbial electrochemical system ran 20 h *prior* to the experiment to reach an equilibrium state. The potential was scanned from -0.7 V to -1.15 V vs Ag/AgCl (3M KCl) at a scan rate of 25 mV s^-1^.

### Limitations of the study

Microbial electrosynthesis confronts a hurdle in achieving industrial viability, primarily attributed to its low EE. Despite achieving a CH_4_-CE exceeding 90%, the overall EE remains at a modest 20-35%, which is comparable to other studies^[12, 17, 36, 37]^ but still distant from commercial feasibility. A techno-economic analysis recommends a minimum EE of 50% for industrial applications of microbial electrosynthesis.^[15]^ Furthermore, for abiotic CO_2_ reduction to value-added chemicals, an EE exceeding 60% is preferable.^[38]^ In a hypothetical calculation, assuming the microbial electrochemical system with the platinized titanium catode achieved a consistent EE of 50%, the maximum volumetric CH_4_ production rate observed on day 12 would need a cell voltage of 2.13 V instead of the 3 V that was applied during the experiment (**Figure S12**). This suggests that a cell voltage of around 2 V or lower would be essential for the optimal operation of the system in question. Optimizing the microbial electrochemical system design, including reducing the anode-cathode distance in a zero-gap configuration using a membrane electrode assembly, is imperative to attain this target.^[12, 13, 14]^ Furthermore, H_2_- and CH_4_-CE fluctuations during experiments underscore the need for a more leak-proof reactor design to minimize losses and enhance H_2_ conversion to CH_4_. This is a general problem when using lab-scale setups and would be mitigated using a large steel bioreactor for industrial applications.

The nickel experiment in this study reveals an average H_2_ volumetric percentage of 32.5% in the headspace, indicating low mass transfer and a loss that needs to be addressed. To enhance soluble H_2_ and align with the microbial electrosynthesis concept of producing H_2_ directly assimilable by microbes, a well-designed bioreactor setup is necessary to prolong H_2_ retention before outgassing. Furthermore, the increased H_2_ bubble formation with rising current density (target of >50 mAcm^-2^) will not allow microbes to directly uptake H_2_ unless the cell density is high enough. Two effective strategies for optimizing H_2_ conversion to CH_4_ involve *either* pressurizing the bioreactor *or* recirculating the off-gas. Both of which should be implemented in the future.

Because heating incurs considerable costs, especially when scaling up for industrial purposes, utilizing the unique heat production capabilities of *M. thermautotrophicus*, which is a hydrogenotrophic methanogen with entropy-retarded growth, is crucial.^[39]^ During methanogenesis, *M. thermautotrophicus* produces heat (negative enthalpy) to offset entropy reduction by producing 1 mole of CH_4_ and 2 moles of water from 5 moles of gas. Unfortunately, exploiting this metabolism advantage for a lab-scale microbial electrosynthesis system is currently unattainable and is achievable only at higher productivities (current densities) and at a lower surface-to-volume ratio.

Another limitation is the occurrence of a pH gradient. The anolyte pH was around 2 for all three experiments, while the catholyte ranged between 7 and 8.5 outside of the shutdown periods. When both chambers have equal pHs, the thermodynamic potential difference between the O_2_ and H_2_ evolution reactions is 1.23 V.^[40]^ In this study, we had a pH gradient that increased the thermodynamic cell voltage to 1.62 V, which is a 0.39 V increase to equal pH conditions. It has been shown that using a vapor-fed anode reduces the pH gradient by not allowing the accumulation of protons.^[13, 40]^ Ideally, the protons will migrate to the cathode chamber, and the large pH difference across the cell will be mitigated. Consequently, a new electrochemical cell, such as a zero-gap electrochemical cell, should be used to optimize the system.^[14]^ Finally, the ion-exchange membrane should be upgraded to avoid gas and water crossovers.

## Conclusion

We observed that intermittent power shutdowns initially affected the microbes, but they quickly adapted to the new experimental conditions. High absolute currents were attained using three electrode materials (*i.e*., platinized titanium, graphite, and nickel), resulting in elevated volumetric CH_4_ production rates. In an extended microbial electrochemical system experiment with nickel as the cathode, incorporating three shutdown types showed minimal influence on overall performance. Specifically, only the short-circuit mode led to higher nickel dissolution, potentially causing increased degradation if employed as the sole shutdown approach. The open-circuit shutdown method stands out as the optimal strategy for system preservation, offering a straightforward approach with minimal disruptive effects, and is recommended as the primary shutdown. To enhance future studies, addressing the influences of O_2_ and CO_2_ crossover is crucial by optimizing both the cell and the recirculating vessel. The platinized titanium cathode outperformed nickel and graphite cathodes. Because of the expensive cost of platinum and possible sulfide contamination throughout the operating period, nickel-based cathodes could be a better option. Especially when nickel-based catalysts, such as NiMo, outperform platinum.^[41]^ Therefore, nickel-based cathodes should be further investigated, emphasizing changing their morphology. We show here that during the testing of new cathodes, shutdown experiments should be performed to also evaluate the effects of industrial casualties such as forced breaks in the operating periods.

## Supporting Information

Supporting information is available online or from the author.

## Acknowledgements

This work was supported by the Deutsche Forschungsgemeinschaft (DFG, German Research Foundation, EBiotech, SPP2240; L.T.A.) – project number 445506379, the Alexander von Humboldt Foundation in the framework of the Alexander von Humboldt Professorship (L.T.A.), and The Novo Nordisk Foundation CO_2_ Research Center with grant number NNF21SA0072700 (L.T.A.). Authors acknowledge funding from the central innovation program for small and medium-sized enterprises (Zentrales Innovationsprogramm Mittelstand [ZIM]), a cooperation project of the ministry of economic affairs and energy (Bundesministeriums für Wirtschaft und Energie [BMWi]) project number 16KN066610 (D.H. and J.R.). This research was also supported by the German Central Innovation Program for small and medium-sized enterprises (SMEs), German Federal Ministry of Education and Research (KMU-innovativ -KMUi-BÖ02: PtGMEC, Förderkennzeichen: 031B1244). The authors gratefully acknowledge the Tübingen Structural Microscopy Core Facility (Funded by the Federal Ministry of Education and Research (BMBF) and the Baden-Württemberg Ministry of Science as part of the Excellence Strategy of the German Federal and State Governments) for their support and assistance in this work. We thank Christian Hochstetter for helping with Figure 1.

