## Supporting Information for "Performance effects from different shutdown methods of three electrode materials for the power-to-gas application with electromethanogenesis"

[c] Prof. Dr. Ir. Largus T. Angenent
*Cluster of Excellence: EXC 2124: Controlling Microbes to Fight Infection*
*University of Tübingen*
*Auf der Morgenstelle 28, 72076 Tübingen*

[d] Prof. Dr. Ir. Largus T. Angenent
*AG Angenent*
*Max Planck Institute for Biology*
*Max Planck Ring 5, 72076 Tübingen, Germany*

[e] Prof. Dr. Ir. Largus T. Angenent
*Department of Biological and Chemical Engineering*
*Aarhus University*
*Gustav Wieds Vej 10D, 8000 Aarhus C, Denmark*

[f] Prof. Dr. Ir. Largus T. Angenent
*The Novo Nordisk Foundation $CO_2$ Research Center (CORC)*
*Aarhus University*
*Gustav Wieds Vej 10C, 8000 Aarhus C, Denmark*

Supporting information for this article is given via a link at the end of the document.

†These authors contributed equally to this work and share first authorship

*Corresponding author

### Supporting Information:

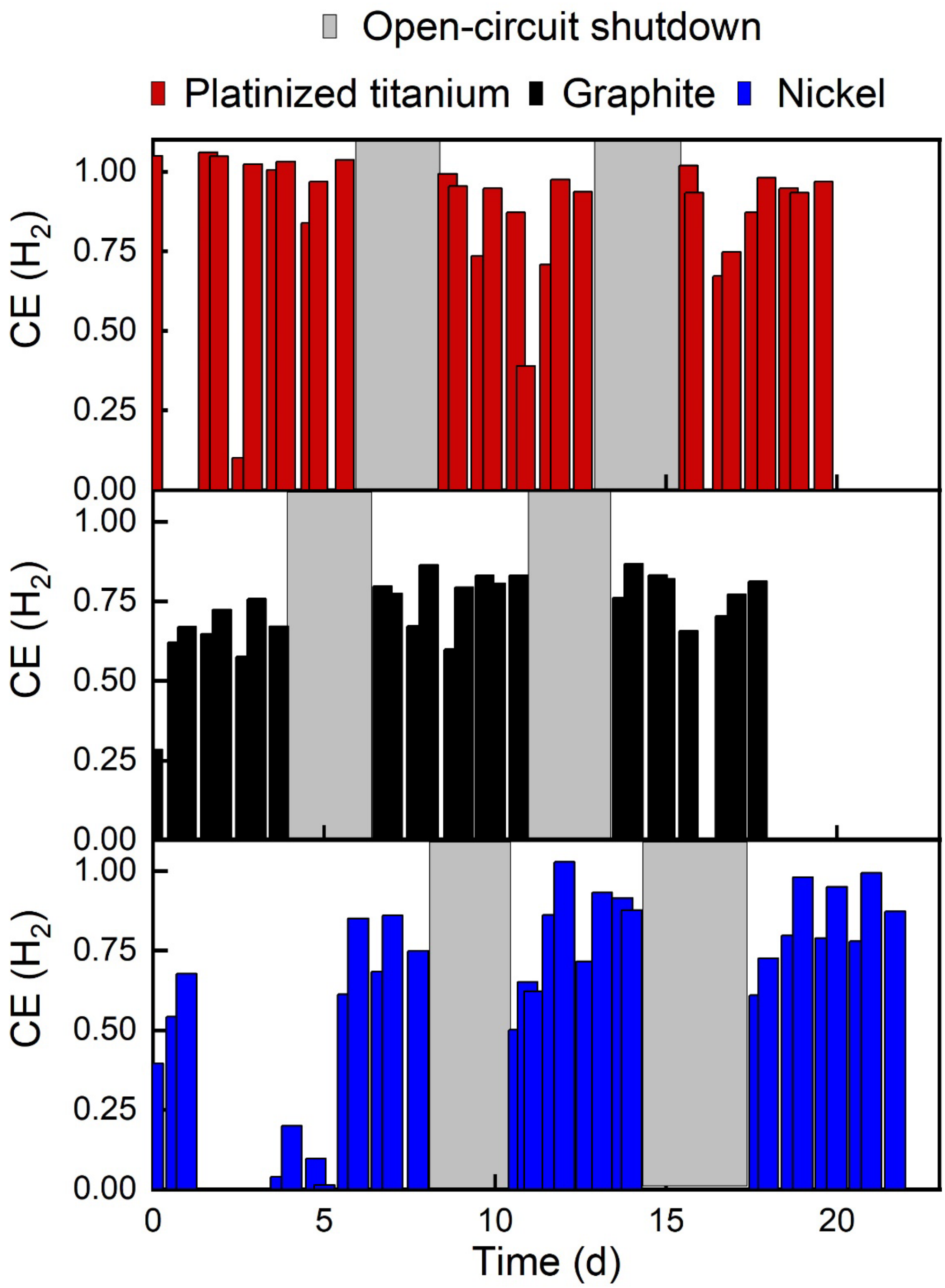


**Figure S1**: CE of $H_2$ during the microbial electrochemical system experiments using platinized titanium, graphite, and nickel as cathode material. The light grey areas correspond to the open-circuit shutdown periods.

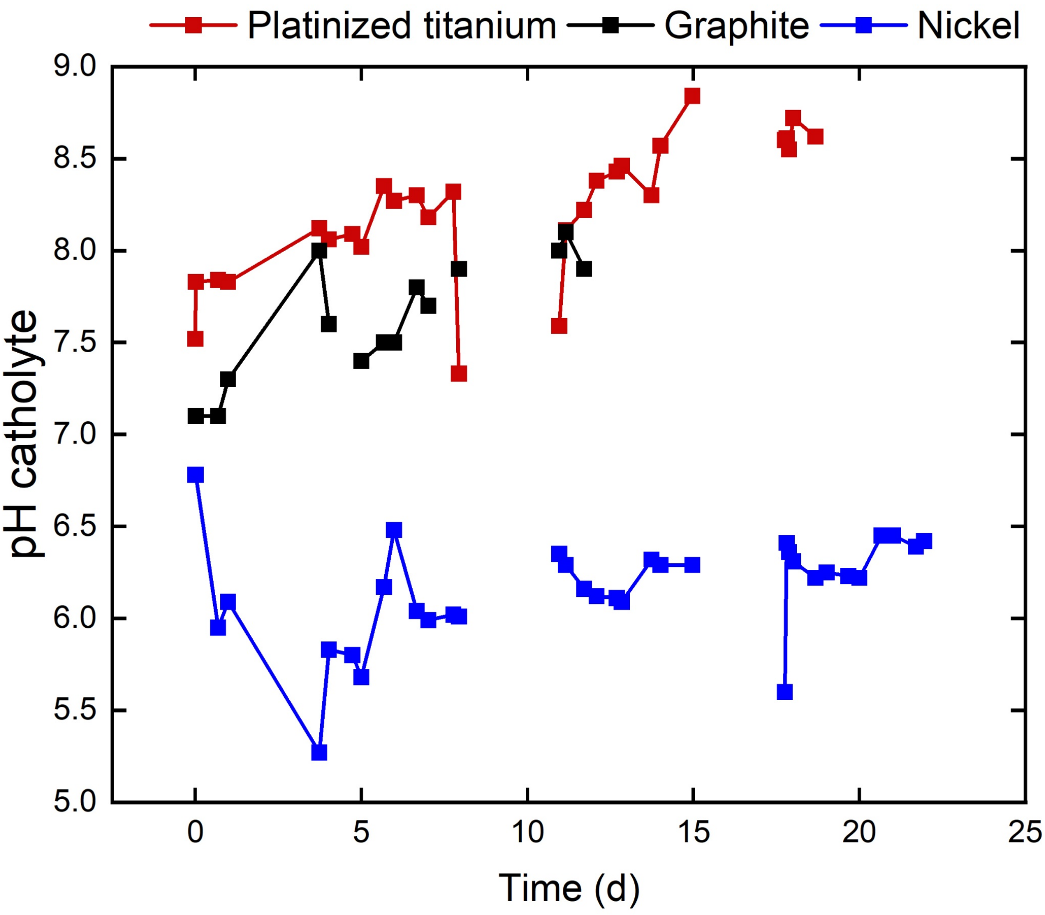


**Figure S2**: catholyte pH of microbial electrochemical system experiments using platinized titanium, graphite, and nickel as cathode material. Interruptions in the line segments correspond to the open-circuit shutdown periods.

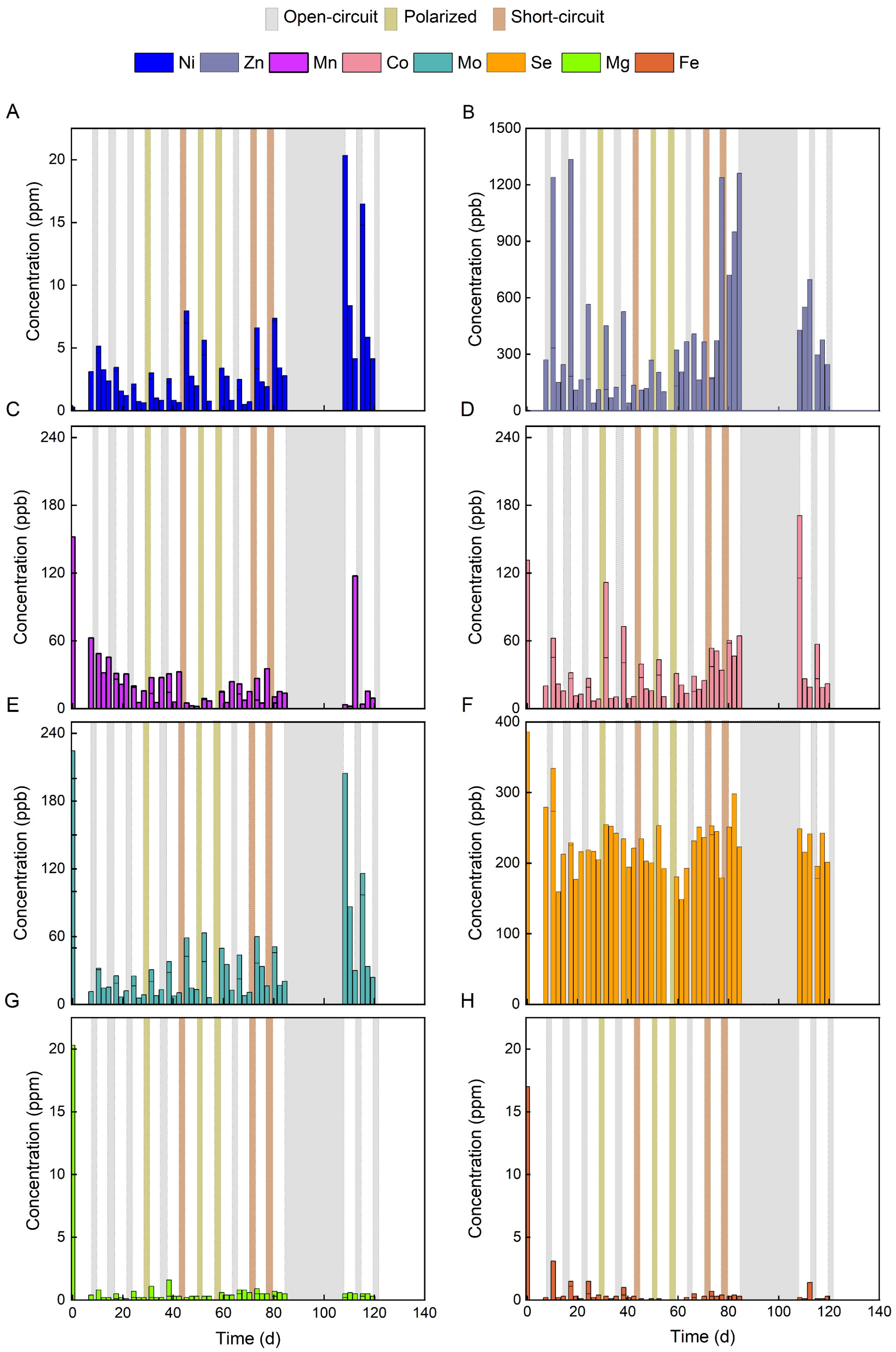


**Figure S3**: Element concentrations of Ni, Zn, Mn, Co, Mo, Se, Mg, and Fe (**A** to **H**) in the catholyte measured with ICP-MS during the microbial electrochemical system with the nickel cathode experiment. Samples were taken at the start, 2 hours, 3 days, and 5 days each week after and before a shutdown procedure. The different shutdown methods are coded in grey as open-circuit shutdown, light yellow as polarized shutdown, and light red as short-circuit shutdown.

A

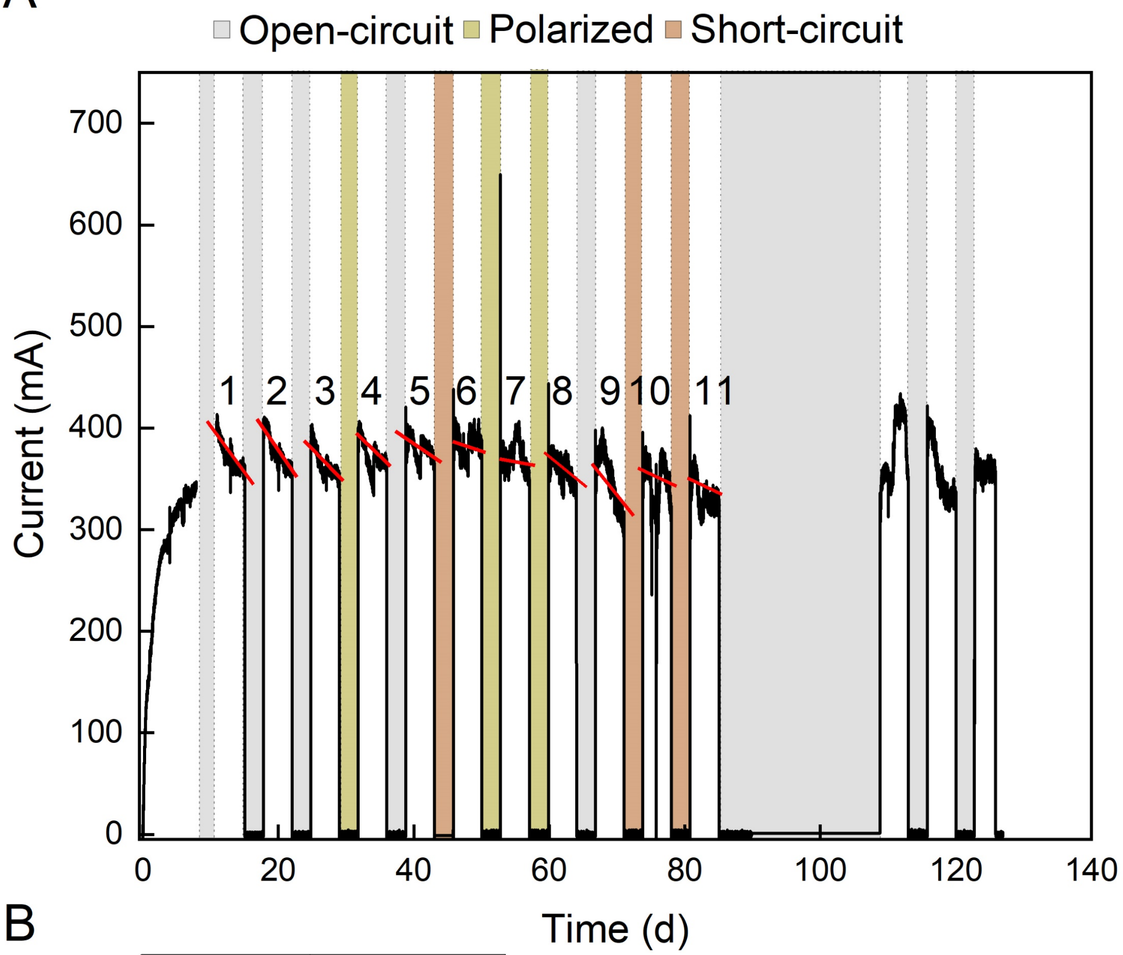


B

| Number | Regression |
|---|---|
| | mA /d |
| 1 | -13.3 |
| 2 | -10.5 |
| 3 | -14.1 |
| 4 | -11.1 |
| 5 | -7.8 |
| 6 | -5.7 |
| 7 | -1.1 |
| 8 | -4.4 |
| 9 | -14.1 |
| 10 | -7.7 |
| 11 | -2.7 |

**Figure S4**: (A) absolute current and its regression (red line) of the microbial electrochemical system experiment using nickel as cathode with different shutdown methods coded in grey as open-circuit shutdown, light yellow as polarized shutdown, and light red as short-circuit shutdown; and (B) the regression analysis of the regression from (A).

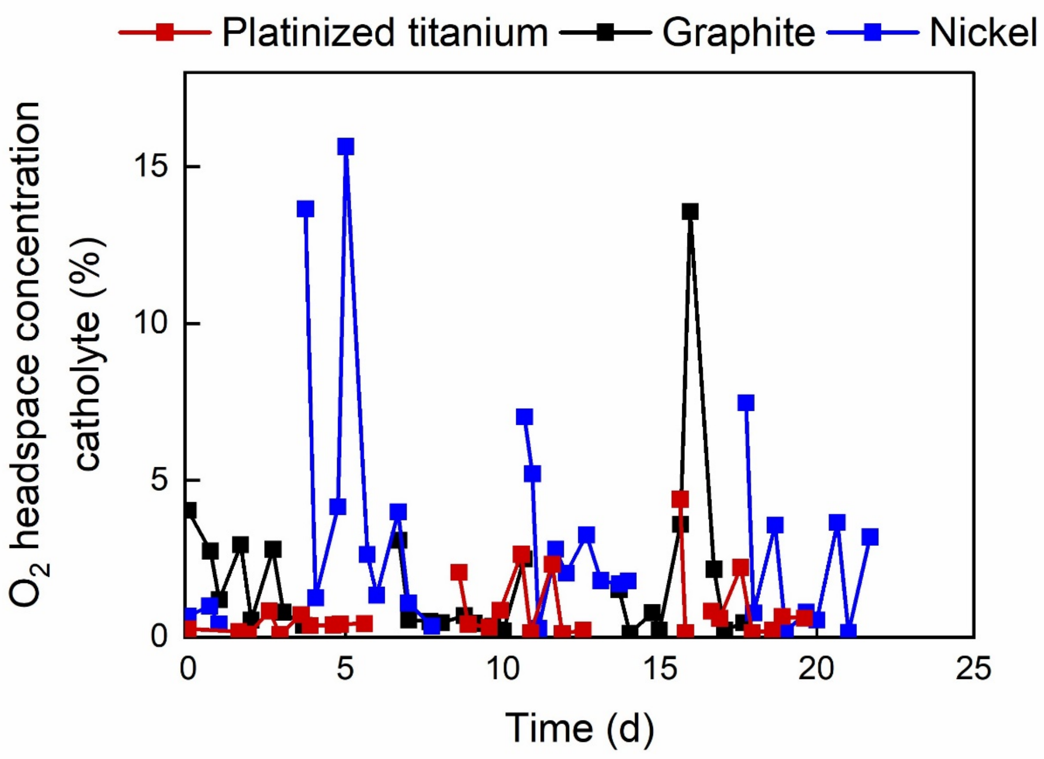


**Figure S5:** The percentage of cathode chamber headspace $O_2$ concentration during the microbial electrochemical system experiment using platinized titanium, graphite, and nickel cathodes.

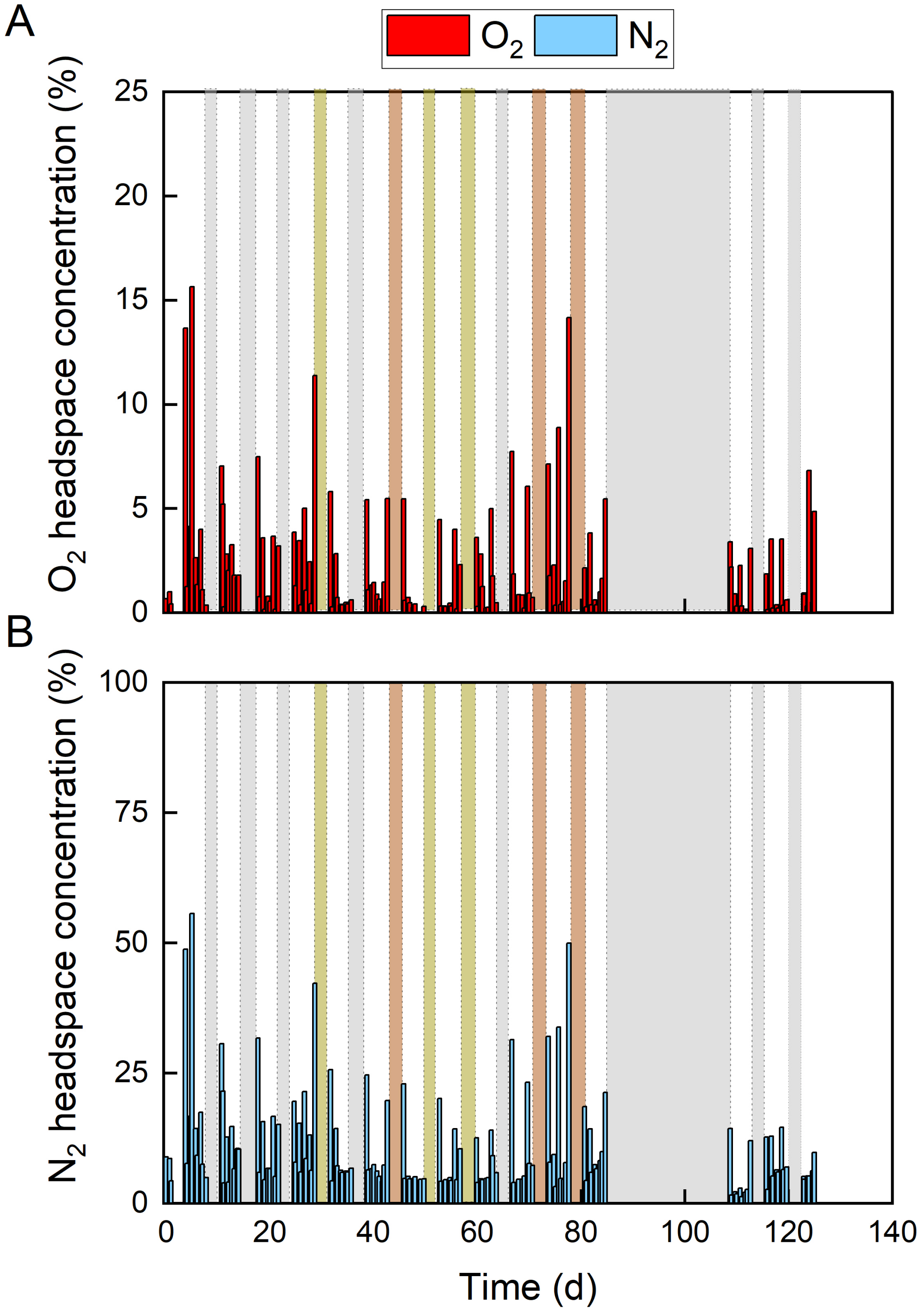


**Figure S6**: cathode chamber headspace concentration in percentage of: (A) $O_2$; and (B) $N_2$ during the microbial electrochemical system experiment using nickel as the cathode. The different shutdown methods are coded in grey as open-circuit shutdown, light yellow as polarized shutdown, and light red as short-circuit shutdown.

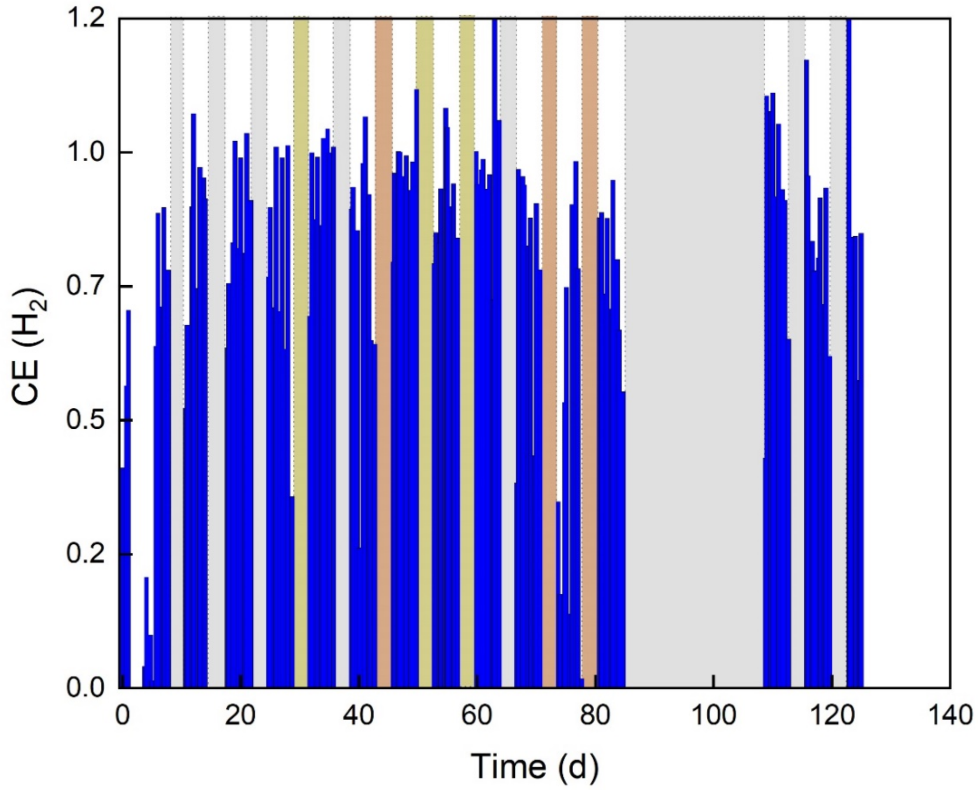


**Figure S7**: Coulombic efficiency (CE) of $H_2$ during the microbial electrochemical system experiment using nickel as the cathode. The different shutdown methods are coded in grey as open-circuit shutdown, light yellow as polarized shutdown, and light red as short-circuit shutdown.

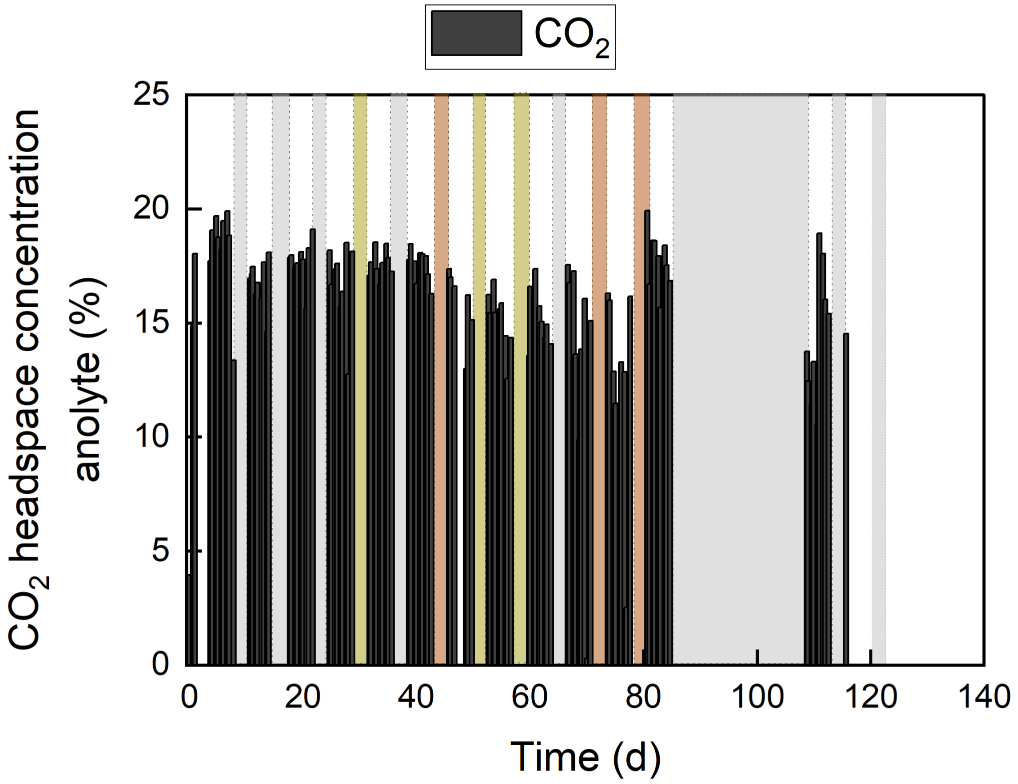


**Figure S8**: anode chamber headspace concentration in % of $CO_2$ during the microbial electrochemical system experiment using nickel as the cathode. The different shutdown methods are coded in grey as open-circuit shutdown, light yellow as polarized shutdown, and light red as short-circuit shutdown.

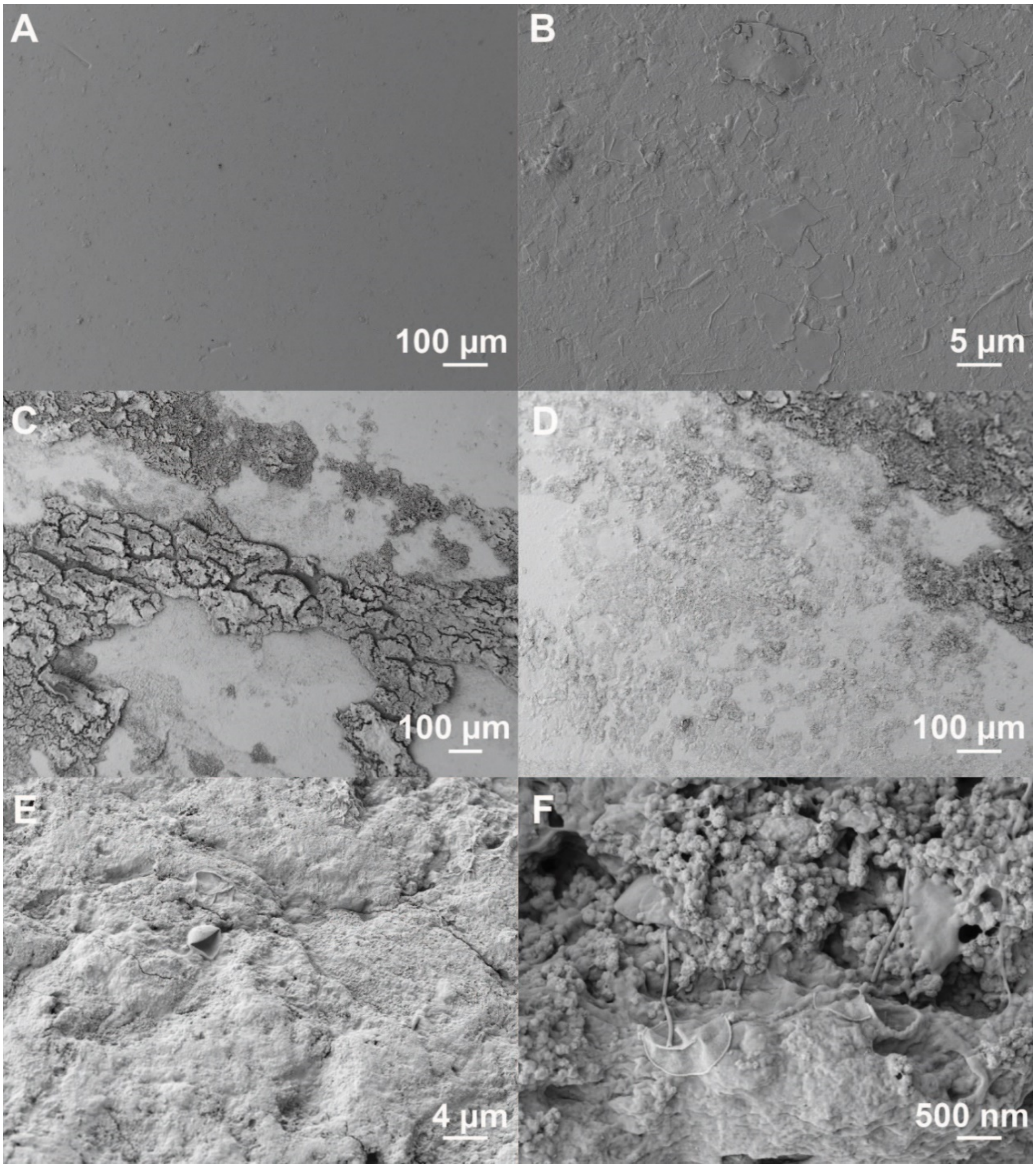


**Figure S9**: SEM micrographs of the: (A) clean Nafion 117 ion-exchange membrane at 100x magnification; (B) close-up of the clean Nafion 117 ion-exchange membrane at 2020x magnification; (C) and (D) the Nafion 117 ion-exchange membrane post-experiment with precipitate at 100x magnification; (E) and (F) close-up of the Nafion 117 ion-exchange membrane post-experiment at 2260x and 20030x magnification, respectively.

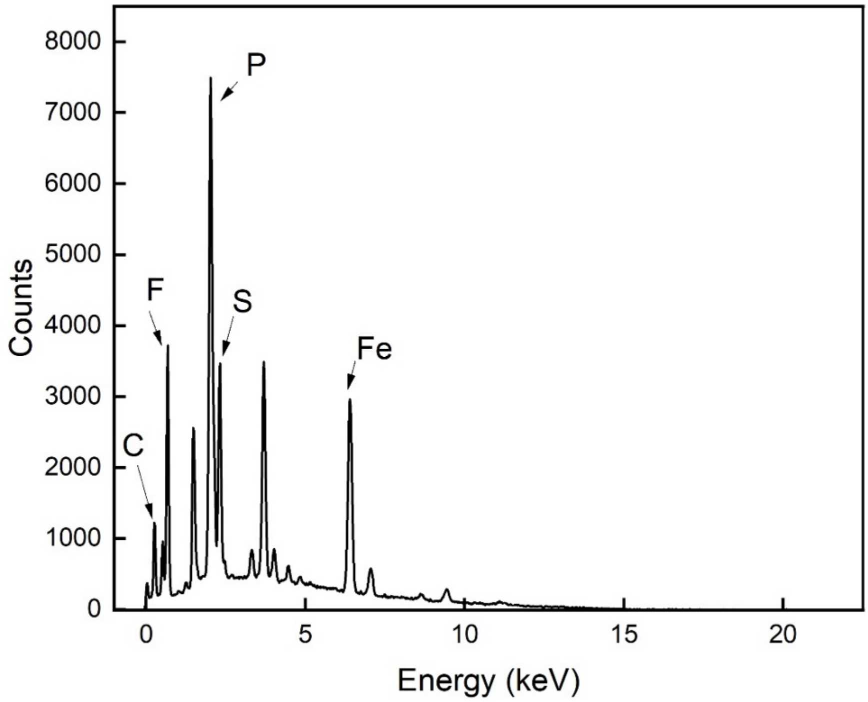


**Figure S10**: EDX Kα elemental mapping spectrum of the Nafion 117 ion-exchange membrane post-experiment.

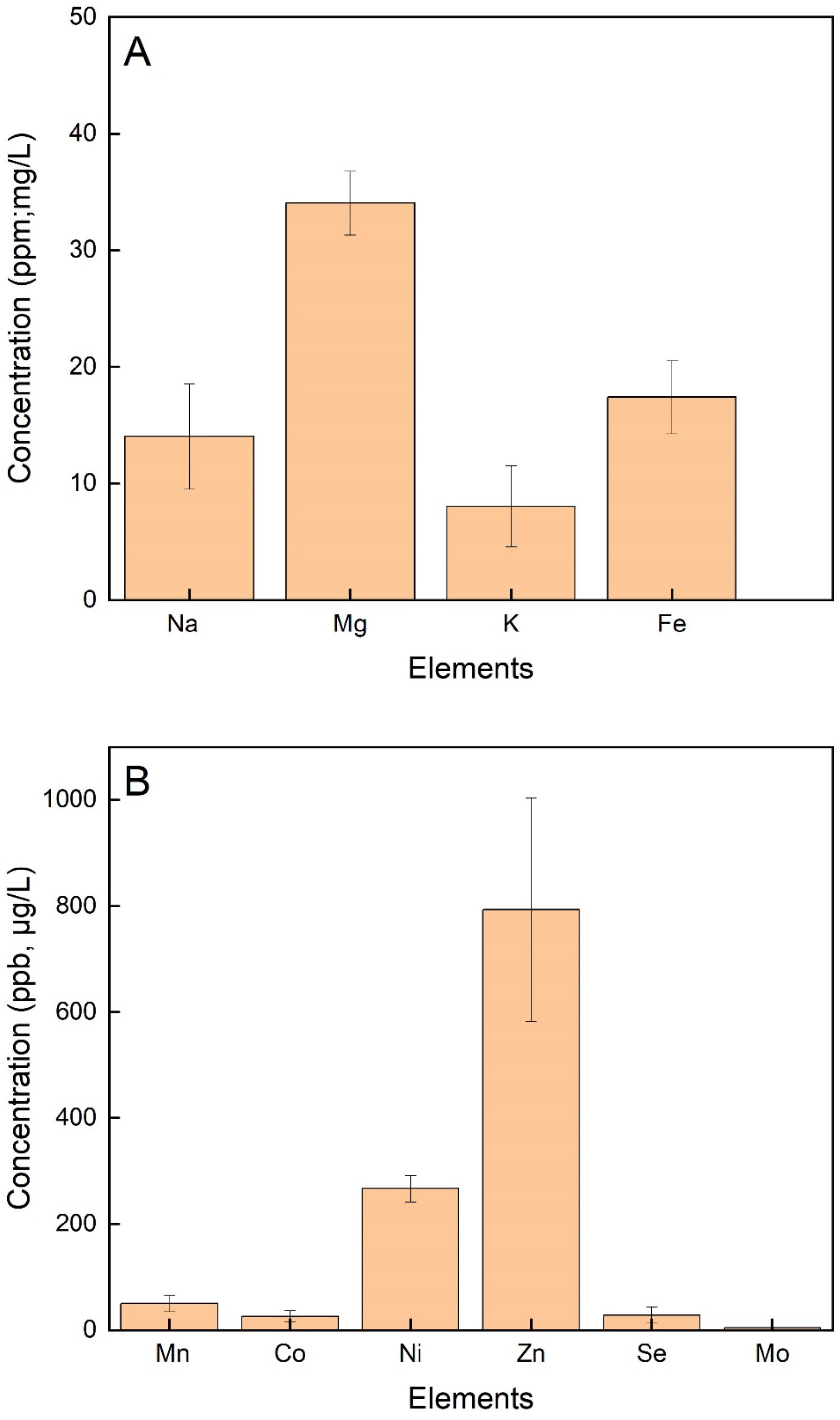


**Figure S11**: Element concentrations of: (**A**) Na, Mg, K, and Fe; and (**B**) Mn, Co, Ni, Zn, Se, and Mo from a 1 $cm^2$ piece of Nafion 117 ion-exchange membrane calcinated. The sample was measured with ICP-MS.

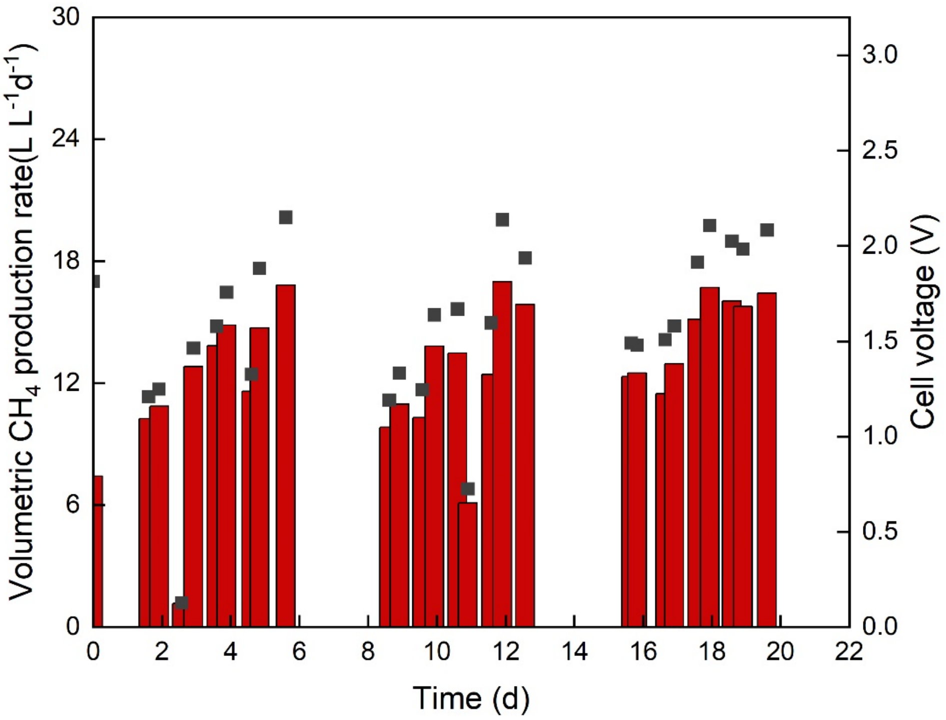


**Figure S12**: Volumetric production rate during the microbial electrochemical system with platinized titanium cathode experiment and its hypothetical cell voltage if a minimum of 50% EE would be reached.
